# A Coarse-Grained Molecular Dynamics Investigation on Spontaneous Binding of Aβ_1-40_ Fibrils with Cholesterol-mixed DPPC Bilayers

**DOI:** 10.1101/2022.09.16.508209

**Authors:** Nikhil Agrawal, Adam A Skelton, Emilio Parisini

## Abstract

Alzheimer’s disease is the most common form of dementia. Its aetiology is characterized by the misfolding and aggregation of amyloid-β (Aβ) peptides into β-sheet-rich Aβ oligomers/fibrils. Whereas experimental studies have suggested that Aβ oligomers/fibrils interact with the cell membranes and perturb their structures and dynamics, the molecular mechanism of this interaction is still not fully understood. In the present work, we have performed a total of 120 μs-long simulations to investigate the interaction between trimeric or hexameric Aβ_1-40_ fibrils with either a 100% DPPC bilayer, a 70% DPPC-30% cholesterol bilayer or a 50% DPPC-50 % cholesterol bilayer. Our simulation data capture the spontaneous binding of the aqueous Aβ_1-40_ fibrils with the membranes and show that the central hydrophobic amino acid cluster, the lysine residue adjacent to it and the C-terminal hydrophobic residues are all involved in the process. Moreover, our data show that while the Aβ_1-40_ fibril does not bind to the 100% DPPC bilayer, its binding affinity for the membrane increases with the amount of cholesterol. Overall, our data suggest that two clusters of hydrophobic residues and one lysine help Aβ_1-40_ fibrils establish stable interactions with a cholesterol-rich DPPC bilayer. These residues are likely to represent potential target regions for the design of inhibitors, thus opening new avenues in structure-based drug design against Aβ oligomer/fibril-membrane interaction.

## 1. Introduction

Amyloid fibrils are misfolded, β-sheet-rich, aggregated proteins that play a key role in more than 20 diseases, including Alzheimer’s disease (AD), Parkinson’s disease (PD), type 2 diabetes, HIV, and different forms of systemic amyloidosis [1-7]. Conditions involving amyloid fibrils formation, which are commonly known as protein misfolding diseases, affect millions of people around the world [8, 9]. According to the amyloid cascade hypothesis, in the brain of AD patients the amyloid β peptide (Aβ) undergoes conformational changes to form water-insoluble Aβ fibrils [10, 11]. These Aβ fibrils then form extracellular neuronal plaques, which represent the major pathological hallmark of AD [12, 13].

Several experimental studies have been performed to reveal the effect of the interaction between Aβ fibrils and lipids/membranes. Han *et al*. used electron tomography to visualize the interaction of Aβ fibrils with lipids of different sizes, also reporting that intracellular fibrils deform the structure of intracellular lipid vesicles and puncture through the vesicular membrane into the cytoplasm [14]. Using simultaneous coherent anti-Stokes Raman scattering (CARS) and 2-photon fluorescence microscopy, Kiskis *et al*. showed that lipids co-localize with fibrillar β-amyloid (Aβ) plaques [15]. In another experimental study, Burns *et al*. revealed the co-localization of cholesterol in Aβ plaques [16]. Previously, several MD simulations studies were also done to investigate the interaction between membranes and Aβ oligomers/fibrils. For instance, Yu *et al*. performed MD simulations of the interaction between Aβ_17-42_ fibrils and a mixed anionic POPC–POPG bilayer [17]. Their study showed that anionic lipids play an important role in the absorption of the Aβ_17-42_ pentamer in the membrane. Tofoleanu *et al*. conducted MD simulations of the interaction between Aβ fibrils and a POPE lipid bilayer, and their data revealed that the charged residues Glu22, Asp23 and Lys28 form electrostatic interactions with head group atoms of the lipid [18]. In another study, Tofoleanu *et al*. used MD simulations to investigate the behaviour of Aβ fibrils with POPC and POPE bilayers [19]. This study revealed that Aβ fibrils form short-lived contacts with POPC head groups and strong contacts with POPE head groups. Dong *et al*. performed MD simulations of Aβ_40_ fibril trimers with a POPG bilayer, revealing that their interaction is mediated by the N-terminal β-sheet [20]. In a recent work, Dias *et al*. used coarse-grained (CG) simulations to investigate the binding of Aβ_1-42_ fibrils with a cholesterol-rich phosphatidylcholine (PC) bilayer, showing that 30% cholesterol is optimal for Aβ_1-42_ fibril interaction with the membrane and that increasing the cholesterol content in PC bilayer up to 50% does not increase the binding frequency of Aβ_1-42_ fibrils with the membrane [21].

It is well established that Aβ fibrils are highly polymorphic in nature and that their two isoforms, Aβ_1-40_ and Aβ_1-42_, show different structural features at the molecular level [22]. Moreover, Aβ fibrils structures have been reported to feature some degree of polymorphism within the same isoform; indeed, Meinhardt [23] *et. al*. showed that Aβ_1-40_ fibrils form different structures even within the same sample and under the same conditions. Hence, this highlights the importance of investigating also the interaction of Aβ_1-40_ fibrils with lipid membranes. In the present study, we aimed to capture the spontaneous binding of the Aβ_1-40_ fibril hexamer and trimer with cholesterol-rich dipalmitoyl phosphatidylcholine (DPPC) bilayer. To this end, we performed sixty independent coarse-grained MD simulations to investigate the interaction of either trimeric or hexameric Aβ_1-40_ fibrils with lipid bilayers of different composition (100% DPPC, 70% DPPC-30% cholesterol, and 50% DPPC-50% cholesterol). The total simulation time was 120 μs (2 µs per run). This allowed us to assess how different concentrations of cholesterol affect the spontaneous binding of trimeric and hexameric Aβ_1-40_ fibrils to the membrane, to identify the nature of the amino acids that are involved in the binding and to determine how the binding process affects fibril solvation in the proximity of the bilayer.

## 2. Methods

### 2.1 Structures and force field of Aβfibrils and DPPC-cholesterol membranes

To perform CG molecular dynamics simulations, we used the NMR structure of an hexameric Aβ_9-40_ fibril (PDB id: 2LMN) [24] and the cryo-electron microscopy (cryo-EM) structure of a trimeric Aβ_10-40_ fibril (PDB id: 6W0O) [25] obtained from the cortical tissue of an AD patient. The atomistic structures of both the hexamer and the trimer were converted into a CG model (Figure 1) using the CHARMM-GUI Martini maker [26, 27] and the N-terminal residues that were missing in both models were added using the CHARMM-GUI model missing residues option. Membrane bilayers of different compositions (100% DPPC, 70% DPPC-30% cholesterol and 50% DPPC-50% cholesterol) were prepared using the CHARMM-GUI membrane bilayer builder option. Table 1 shows the initial number of lipid molecules in each type of membrane. Prior to starting the simulations on the fibril-membrane complexes, membranes were equilibrated for 200 ns each and the average area per lipid was calculated for the last 50 ns for each simulation (Supplementary Table 1).

**Table1.** Number of lipids in each membrane bilayers.

| Membrane Bilayer | Number of DPPC molecules | Number of cholesterol molecules |
| --- | --- | --- |
| 100 % DPPC | 800 (400 in each leaflet) | 0 |
| 70% DPPC-30% cholesterol | 560 (270 in each leaflet) | 240 (120 in each leaflet) |
| 50% DPPC-30% cholesterol | 400 (200 in each leaflet) | 400 (200 in each leaflet) |

**Figure 1.**
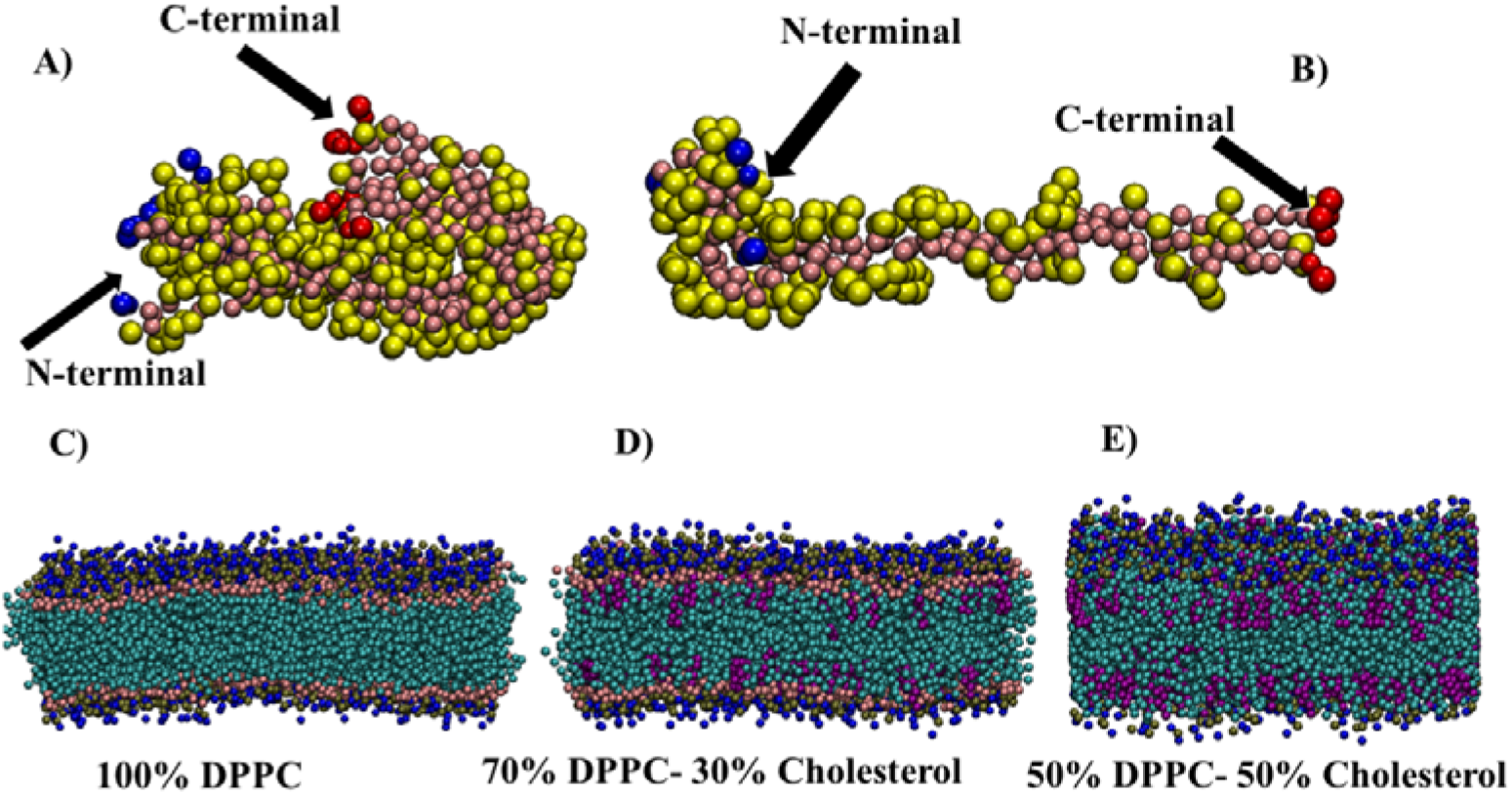
A) CG model of the Aβ_1-40_ fibril hexamer (PDB id: 2LMN). B) CG model of the Aβ_1-40_ fibril trimer (PDB id: 6W0O). C) 100% DPPC membrane bilayer. D) 70% DPPC-30% cholesterol membrane bilayer D) 50% DPPC-50% cholesterol membrane bilayer. CG models of fibrils are shown using VdW representation, while backbone beads of fibrils are shown in pink colour, side chain beads are shown in yellow colour, N-terminal Asp1 residue is shown in blue colour and C-terminal Val40 is shown in red colour. The head group beads NC3 and PO4 of the DPPC molecules are illustrated in blue and tan, respectively, while the head group of cholesterol molecules is colored maroon. The tails of both lipid molecules are depicted in cyan.

The Martini 2.2 protein force field [28] was used for the Aβ_1-40_ fibrils, while the Martini 2.0 force field was used for the membrane, water and ions [29].

### 2.2 Simulation protocol

We studied a total of six Aβ_1-40_ fibril-membrane systems. System one contains one Aβ_1-40_ fibril hexamer, a 100% DPPC bilayer and 49905 water molecules. System two contains one Aβ_1-40_ fibril hexamer, a 70% DPPC-30% cholesterol bilayer and 37445 water molecules. System three contains one Aβ_1-40_ fibril hexamer, a 50% DPPC-50% cholesterol bilayer and 32478 water molecules. A total of 18 NA^+^ ions were added in these three systems to neutralize their overall charge. System four contains one Aβ_1-40_ fibril trimer, a 100% DPPC bilayer and 50081 water molecules. System five contains one Aβ_1-40_ fibril trimer, a 70% DPPC-30% cholesterol bilayer and 37608 water molecules. System six contains one Aβ_1-40_ fibril trimer, a 50% DPPC-50% cholesterol bilayer and 32699 water molecules. A total of 9 NA^+^ were added to neutralize the overall charge of these three systems. Initially, all six systems were energy-minimized using the steepest descent algorithm [30]. All systems were equilibrated in five cycles, with decreasing restrains in each cycles from force constant 1000 kJ mol^−1^ nm^−2^ in the first cycle to 50 kJ mol^−1^ nm^−2^ in the last cycle on the Aβ_1-40_ fibril and from 20kJ mol^−1^ nm^−2^ in the first cycle to 10 kJmol^−1^mol^−2^ in the last cycle on the lipid head groups. Equilibration was performed for a total 4750 ps. All production simulations were done without any restrains on the Aβ_1-40_ fibrils and membranes. The Parrinello-Rahman algorithm [31] with semi-isotropic pressure coupling type was used for pressure coupling and the velocity-rescale algorithm [32] was used for temperature coupling. Pressure coupling and temperature bath times were set at 12.0 ps and 1.0 ps, respectively. All simulations were performed at the temperature of 310.15 K and at the pressure of 1 atm. A 30 fs time step was used for the integration of Newton’s equations of motion. A cut-off of 1.1 nm was employed for Van der Waals and electrostatic interactions, and electrostatic interactions were treated using the reaction field method. A total of 60 independent simulations were done using an initial random velocity generated by GROMACS 2021 [33]. Each simulation was performed for 2 µs using GROMACS 2021 simulation package [33].

### 2.3 Analysis details

The minimum distance of the Aβ_1-40_ fibrils residues from the membrane was calculated using the GROMACS minidist program. An interaction with a Aβ_1-40_ fibrils residue was taken into account when the minimum distance between membrane lipids (either DPPC or cholesterol) and the residue was ≤ 5 Å. The minimum distance between each residue and the membrane was calculated in the time range of 0-2 µs. The number of water and lipid molecules was calculated within 6 Å of the Aβ_1-40_ fibrils. Area per lipid was calculated using FATSLiM[34]

## 3. Results

### 3.1 Binding of the A_1-40_ fibrils with the membranes

Cytotoxicity of amyloid fibrils is generally reported to their binding with cell membranes [35, 36]. To see the spontaneous binding of Aβ_1-40_ fibrils (hexamer and trimer) with the membrane, we calculated the time evolution of the minimum distance between hexamer and trimer with all three bilayers (Figure 2). We observed no spontaneous binding in any of the 20 independent trajectories (10 each for the Aβ_1-40_ hexamer and the trimer, Figure 2A, D) with the 100% DPPC bilayer. Indeed, although we noticed a period of time when fibrils came close to DPPC bilayer to establish some transient interactions, the distance between the fibrils and the membrane remained ≥ 5 Å for most of the simulation time. Conversely, out of 20 independent trajectories of the Aβ_1-40_ fibrils with the membrane containing 70% DPPC and 30% cholesterol (Figure 2B, E), in two trajectories of the trimeric Aβ_1-40_ fibrils (SIM3 and SIM10) we observed stable binding with the membrane. In particular, the Aβ_1-40_ trimer was found within 5 Å from the membrane for around ∼0.915 µs and ∼0.575 µs in SIM3 and SIM10, respectively. In both of these simulations, the trimer remained bound to the membrane until the end of the simulation time. In all other trajectories, only transient interactions with the membrane were noted. When considering 50% DPPC-50% cholesterol membranes (Figure 2C, E), out of 10 simulations with hexameric Aβ_1-40_ fibrils, we observed stable binding in 6 trajectories (SIM1, SIM4, SIM7, SIM8, SIM9, and SIM10), while out of 10 trajectories with Aβ_1-40_ trimer, stable binding with the membrane was observed in 5 trajectories (SIM1, SIM4, SIM5, SIM6, SIM9). The longest binding time was detected in SIM1 and SIM4 for the hexamer simulations, and in SIM4 and SIM9 fir the trimer simulations. In all these 4 simulations, Aβ_1-40_ fibrils were found to bind with the membrane for more than 50% of the total simulation time.

**Figure 2.**
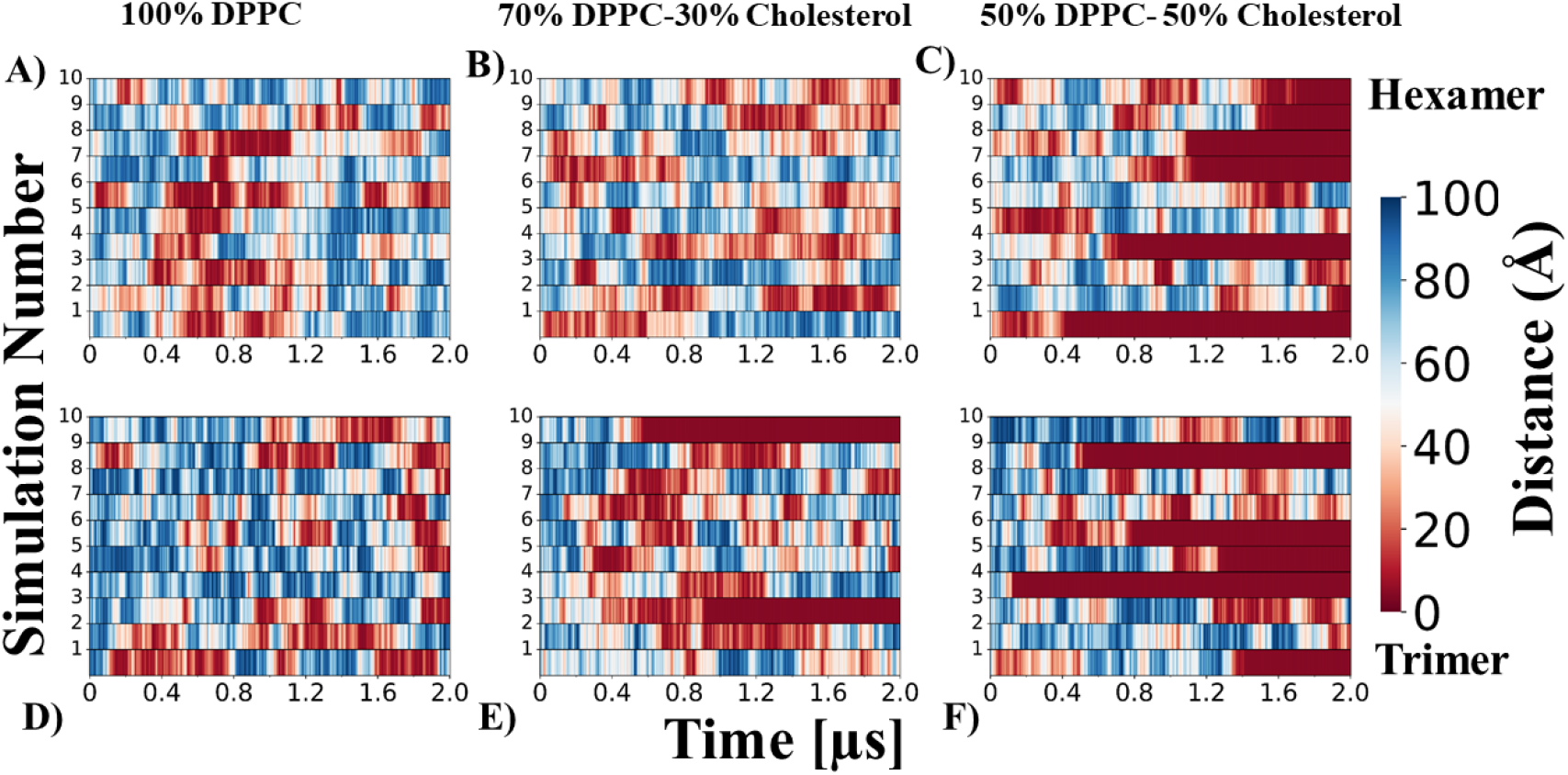
A-C) Time evolution of the minimum distance between Aβ_1-40_ fibrils hexamer and 100% DPPC, 70% DPPC-30% cholesterol and 50% DPPC-50% cholesterol membranes, respectively. D-F) Time evolution of the minimum distance between Aβ_1-40_ fibrils trimer and 100% DPPC, 70% DPPC-30% cholesterol and 50% DPPC-50% cholesterol membranes, respectively.

To further evaluate the distribution of minimum distances in all the simulations, we constructed violin plots for both types of fibrils with all the three different bilayers (Figure 3). The Aβ_1-40_ fibril hexamer violin plot shows high density of values within 5Å for the bilayer containing 50 % cholesterol (Figure 3A, red colour). In the case of trimer violin plot (Figure 3B), the highest density of values within 5Å is also seen for the membrane containing 50% cholesterol (Figure 3B, red colour), followed by the membrane containing 30% cholesterol (Figure 3B, blue colour). Violin plots also reveal a decrease in minimum distance values in going from 0% cholesterol to 50% cholesterol in the membrane for both type of fibrils. Figure 4 shows Aβ_1-40_ fibrils (hexamer and trimer) with membrane containing 50% DPPC and 50% cholesterol at four different time points in two representative trajectories.

**Figure 3.**
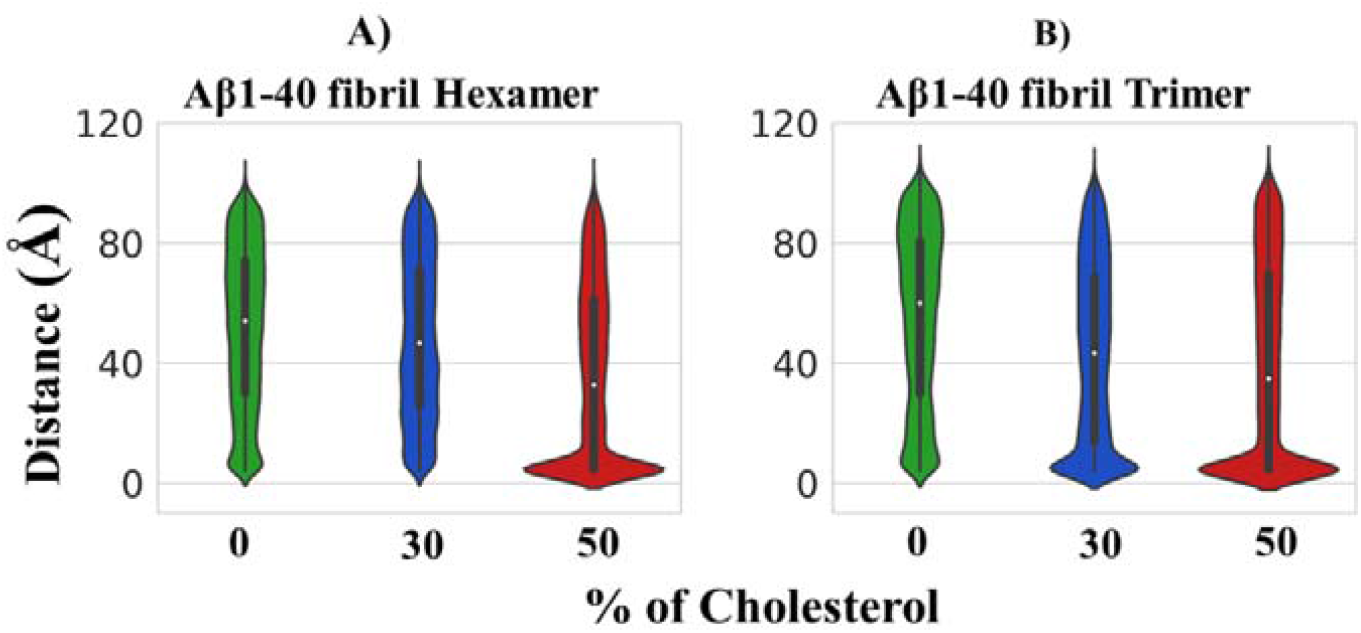
Minimum distance violin plots for different concentrations of cholesterol in the membrane. A) Aβ_1-40_ hexamer, B) Aβ_1-40_ trimer.

**Figure 4.**
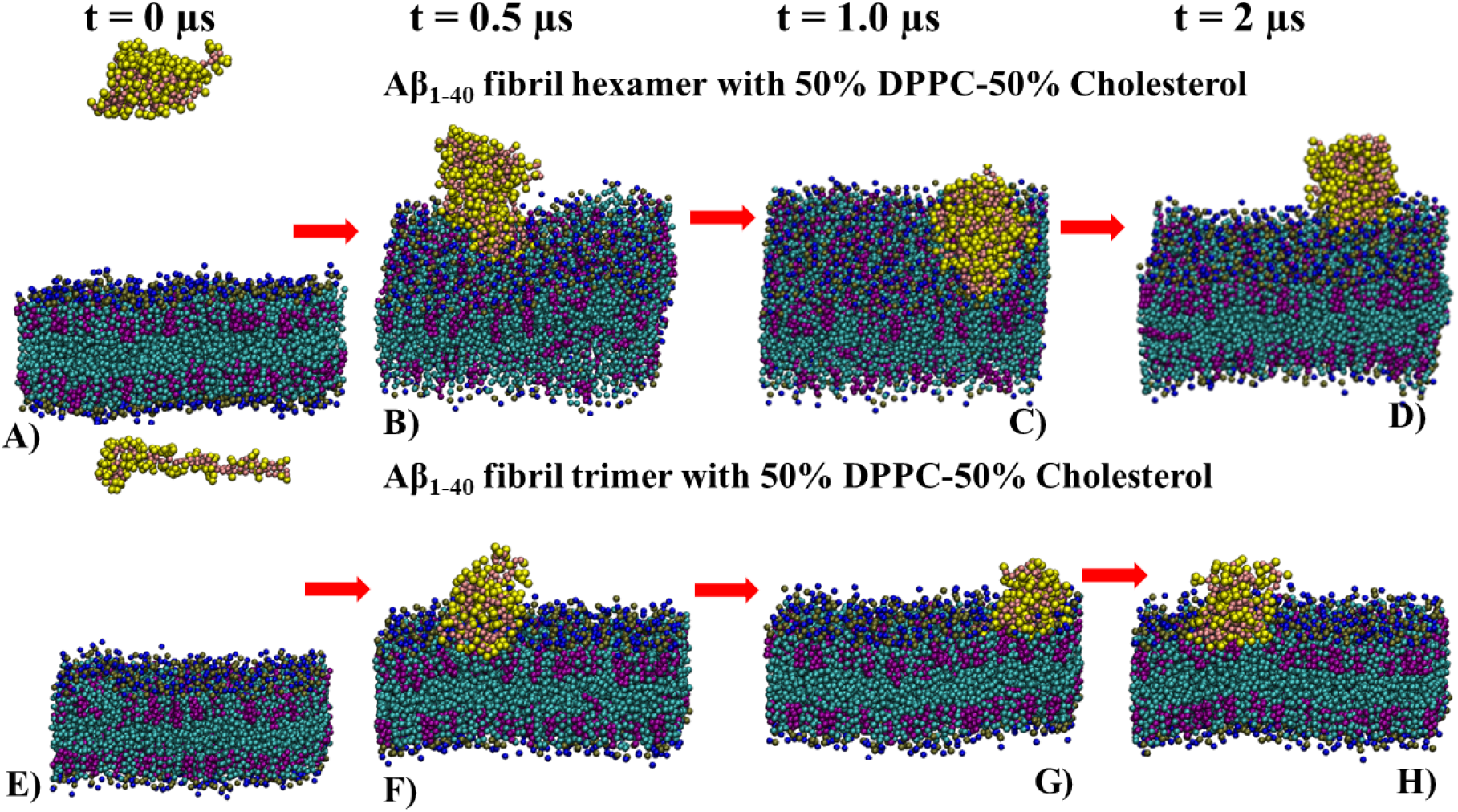
Structures of the Aβ_1-40_ fibrils with membrane containing 50% DPPC and 50% cholesterol at four different time points from two representative trajectories. A-D) Representative image for Aβ_1-40_ fibril trimer at 0, 0.5, 1.0 and 2.0 µs. E-H) Representative image for Aβ_1-40_ fibril hexamer at 0, 0.5, 1.0 and 2.0 µs for Aβ_1-40_ fibril hexamer. Aβ_1-40_ fibrils and membranes are shown in VdW representation. The backbone beads of Aβ_1-40_ fibrils are depicted in pink, and the side chains beads are shown in yellow. The head group beads NC3 and PO4 of the DPPC molecules are illustrated in blue and tan, respectively, while the head group of cholesterol molecules is colored maroon. The tails of both lipid molecules are depicted in cyan.

Overall, these data suggest that, if we exclude some transient interactions, Aβ_1-40_ fibrils (either hexamer or trimer) show little to no affinity for cholesterol-free membranes. However, when the cholesterol level increases to 50% both types of fibrils bind stably to the membrane. Moreover, Aβ_1-40_ fibrils can also bind firmly with the membrane when 30% cholesterol is present, as seen in trimer simulations.

### 3.2 Identification of Membrane Binding residues

To identify which residues bind to the membrane, we calculated the percentage of simulation time for which the minimum distance of each residue from the membrane is ≤ 5Å for more than 5% of the simulation time (Figure 4). In simulations of Aβ_1-40_ fibrils with membrane containing 100% DPPC, no residue binding for more than 5% of the simulation time was observed in all 20 trajectories (supplementary figure 1, 2). In Aβ_1-40_ fibrils simulations with 70% DPPC-30% cholesterol membrane, out of 20 trajectories (10 each for hexamer and trimer), we detected residues of the Aβ_1-40_ fibril trimer at ≤ 5Å for more than 5% of the simulation time in 2 trajectories (SIM3 and SIM10). In both of these simulations, residues from all three chains participated in membrane binding (Figure 5).

**Figure 5.**
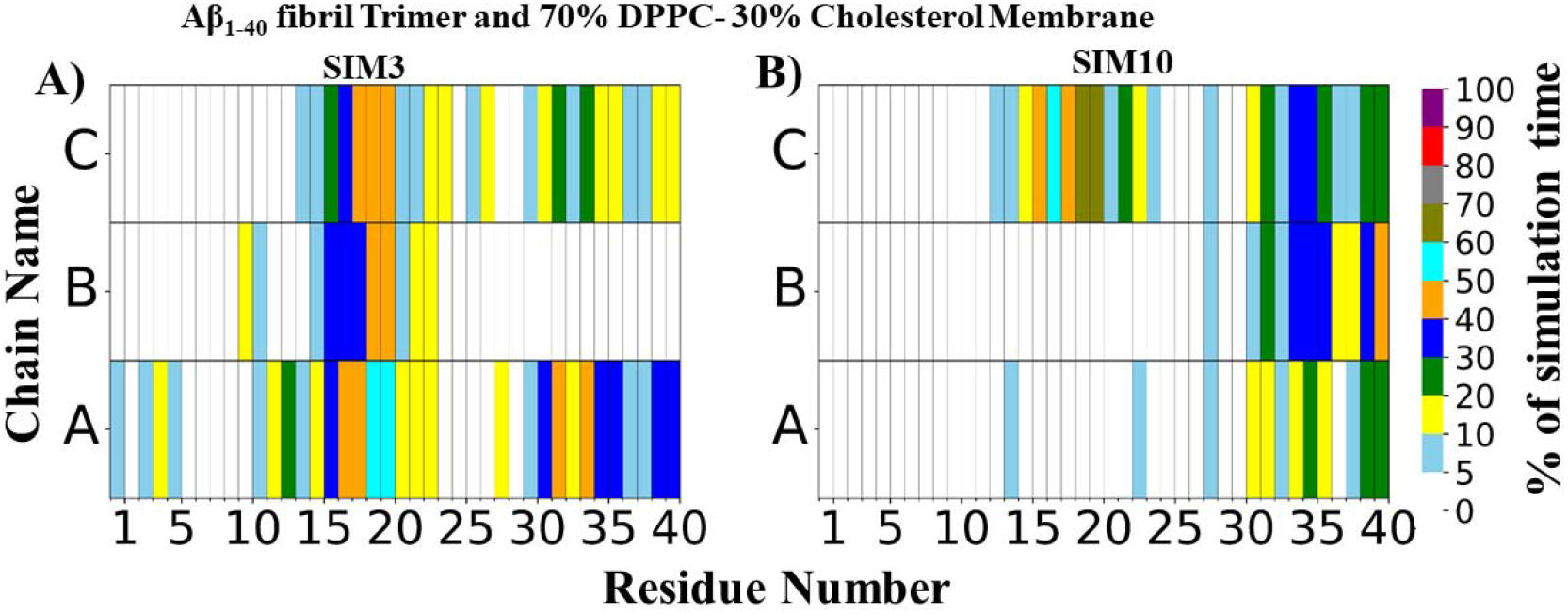
Percentage of time in which residues of the Aβ_1-40_ fibril trimer remained at a distance ≤ 5Å from the 70% DPPC-30% cholesterol membrane in 2 out of 10 trajectories. A) SIM3, B) SIM10.

In both of these simulations, the binding process is mostly mediated by amino acids from the middle and the C-terminal region of the fibrils. Lys16, Leu17, Val18, Phe19, Phe20, Val39, Leu34, Met35, Val36, Val39 and Val40 are the residues that are mostly involved in the binding with the membrane in these two simulations. These residues interact with the membrane for more than 30% of the simulation time.

With a 50% DPPC-50% cholesterol membrane, we observed residues binding to the membrane for more than 5% of the simulation time in 11 trajectories out of 20 (10 hexamers and 10 trimers). In 6 out of 10 hexamer simulations (SIM1, SIM4, SIM7, SIM8, SIM9 and SIM10), several amino acids of the Aβ_1-40_ fibrils were found to bind the membrane (Figure 6). In particular, in SIM1, 3 chains, in SIM5 and SIM7, 5 chains, in SIM8 all 6 chains, and in SIM9 and SIM10, 4 chains participated in the binding process. In all these simulations, the strongest binding was detected for the C-terminal residues (Ile31, Leu34, Met35, Val36, Val39, Val40) and for the middle region residues (Lys16, Leu17, Val18, Phe19, Phe20). Other than these residues, some binding involving N-terminal residues (Asp1, Glu3, Phe4, and Arg5) was also observed for a short period of time (less than 30% of the simulation time) in some of the trajectories.

**Figure 6.**
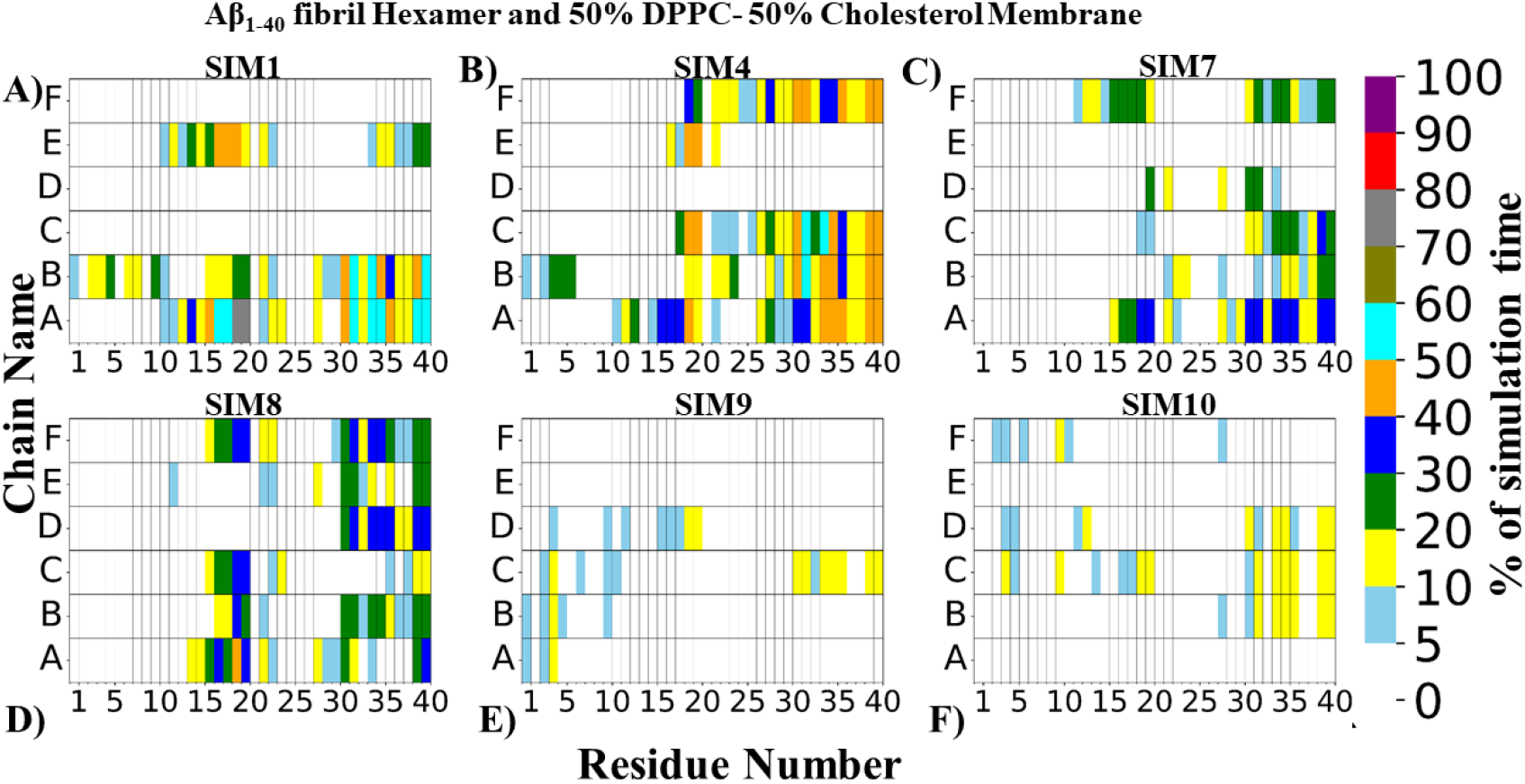
Percentage of time in which residues of the Aβ_1-40_ fibril hexamer remained at a distance ≤ 5Å from the 50% DPPC-50% cholesterol membrane in 6 out of 10 trajectories. A) SIM1, B) SIM4. C) SIM7, D) SIM8, E) SIM9, F) SIM10.

In Aβ_1-40_ fibril trimer simulations, in all five trajectories where residues were found to bind to the membrane (SIM1, SIM4, SIM5, SIM6 and SIM9) (Figure 7), the strongest binding was observed for residues in the middle region (Lys16, Leu17, Val18, Phe19, Phe20) and for residues in the C-terminal region (Ile31, Ile32 Leu34, Met35, Val36, Val39, Val40). For a short period of time, binding was observed also for N-terminal residues (Asp1, Glu3, Phe4, and Arg5).

**Figure 7.**
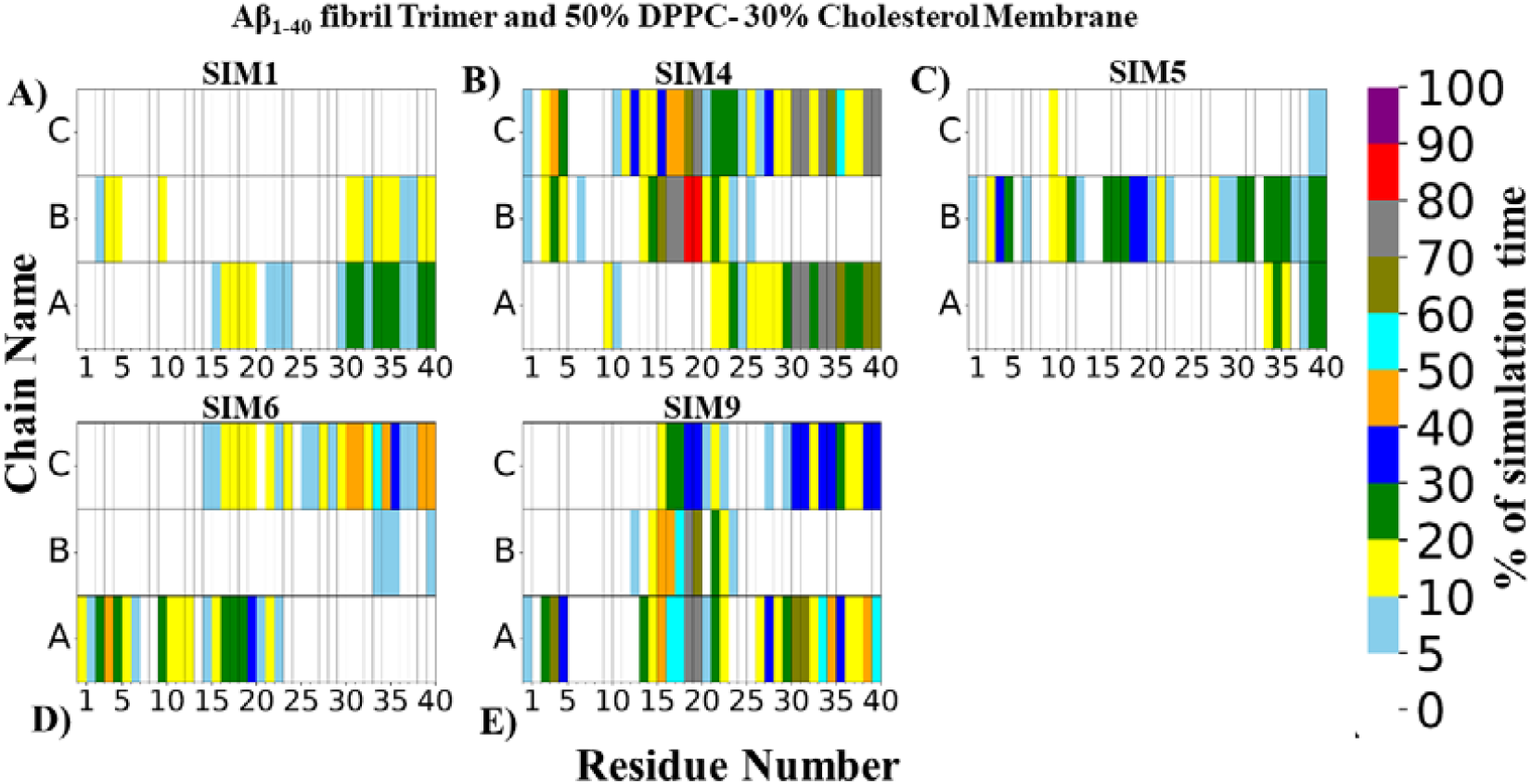
Percentage of time in which residues of the Aβ_1-40_ fibril trimer remained at a distance ≤ 5Å from the 50% DPPC-50% cholesterol membrane in 5 out of 10 trajectories. A) SIM1, B) SIM4, C) SIM5, D) SIM6, E) SIM7.

Overall, the binding to membranes containing either 30% cholesterol or 50% cholesterol involves similar Aβ_1-40_ fibril residues, mostly those that are located in the middle region (the positively charged Lys16, the hydrophobic Leu17 and Val18, the aromatic amino acids Phe19 and Phe20) and those that are found at the C-terminus (the hydrophobic Val36, Val39, Val40, Ile31 and Met35).

### 3.3 Time evolution of the number of water, lipid molecules around Aβ_1-40_ fibrils

To quantify the number of water molecules that the Aβ_1-40_ fibrils must displace to interact with the membrane, we calculated the time evolution of the number of water molecules within 6Å of the Aβ_1-40_ fibrils for all 60 independent trajectories (supplementary figure 3, 4). To interact with the membrane, Aβ_1-40_ fibril hexamer lost ∼150-180 water molecules and Aβ_1- 40_ fibril trimer lost ∼120-130 water molecules. To quantify how many lipid molecules were found in proximity of the Aβ_1-40_ fibrils, we calculated the time evolution of the number of lipid molecules for all 60 trajectories. An almost negligible number of lipids remained within 6Å of Aβ_1-40_ fibrils with 100% DPPC membrane in all 20 trajectories. The same occurred when considering Aβ_1-40_ fibril hexamer with membrane containing 70% DPPC and 30% cholesterol. In the case of Aβ_1-40_ fibril trimer, ∼3-6 cholesterol molecules and ∼6-18 DPPC molecules were found to remain within 6Å from the membrane in the 2 trajectories (SIM3, and SIM10) in which binding occurred (Figure 8).

**Figure 8:**
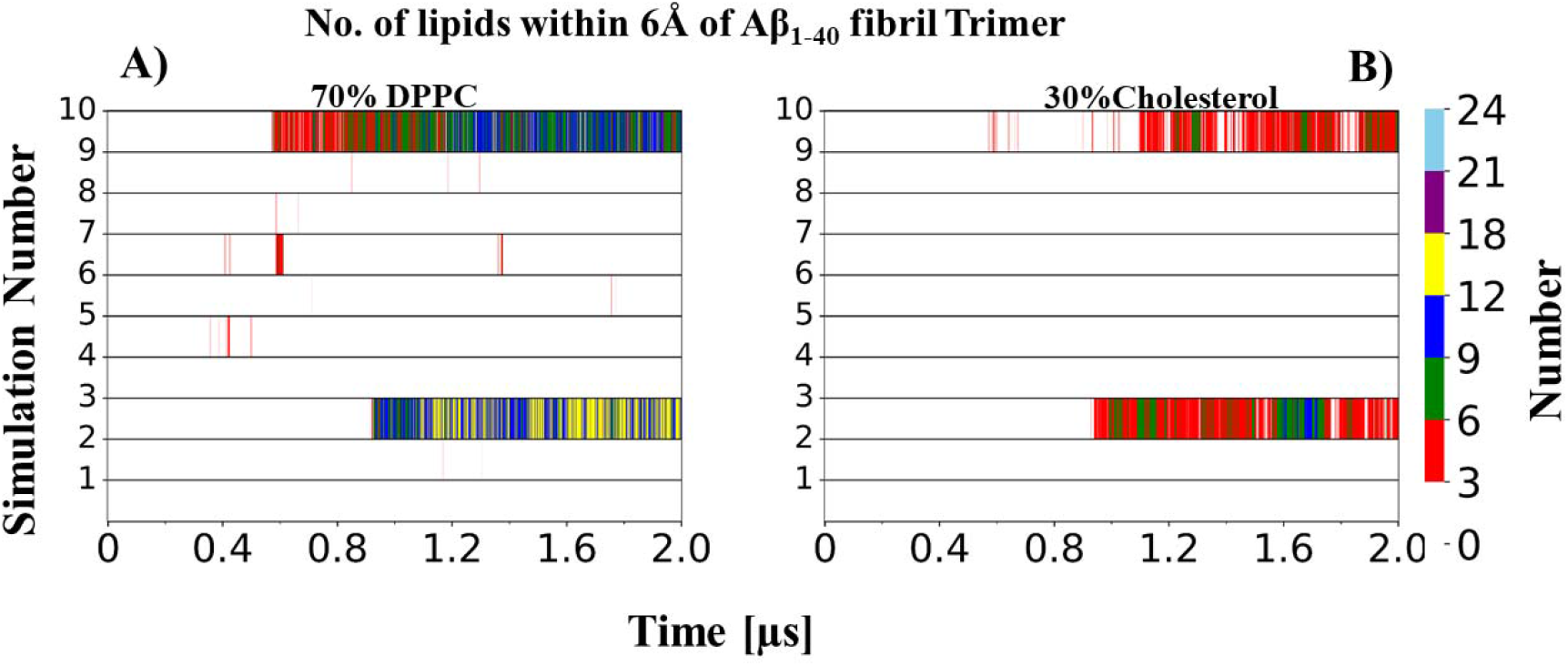
Time evolution of the number of lipids within 6Å of a Aβ_1-40_ fibril trimer from a 70% DPPC-30% cholesterol membrane. A) DPPC, B) Cholesterol

**Figure 9:**
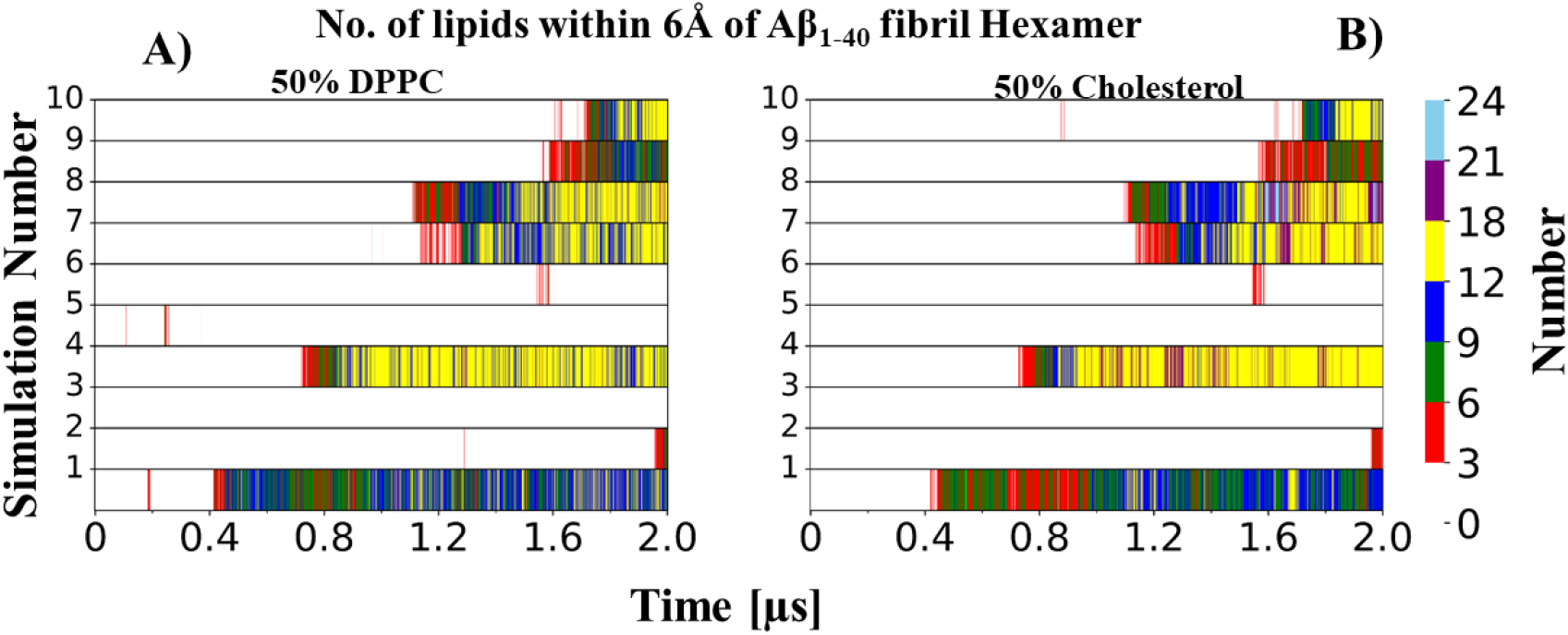
Time evolution of the number of lipids within 6Å of a Aβ_1-40_ fibril hexamer from a 50% DPPC-50% cholesterol membrane. A) DPPC, B) Cholesterol

In simulations of Aβ_1-40_ fibril hexamer with a 50% DPPC-50% cholesterol membrane, we observed an almost equal number of DPPC and cholesterol lipids within 6Å of the Aβ_1-40_ fibril.

In the case of Aβ_1-40_ fibril trimer with membrane containing 50% DPPC and 50% cholesterol, we noticed a generally higher number of cholesterol lipids than DPPC lipids within 6Å of Aβ_1-40_ fibrils (Figure 10, A and B).

**Figure 10:**
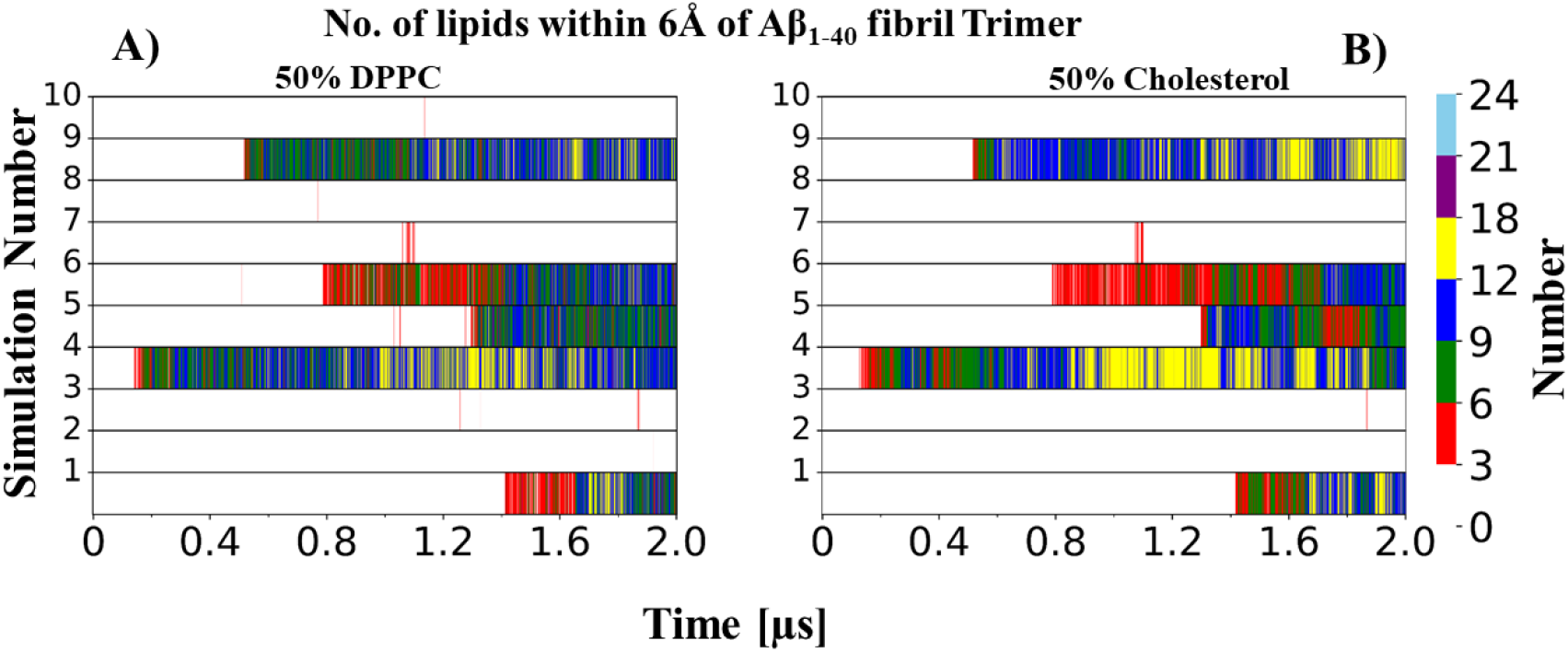
Time evolution of the number of lipids within 6Å of a Aβ_1-40_ fibril trimer with a 50% DPPC-50% cholesterol membrane. A) DPPC, B) Cholesterol.

Overall, these data suggest that an increase in the percentage of cholesterol in the membrane leads to a greater number of cholesterol lipids within 6Å of the fibrils. This could be a possible reason for Aβ fibrils to bind more firmly to the membrane when the percentages of cholesterol and DPPC are similar.

## 4. Discussion

In this study, we performed coarse-grained MD simulations of the interaction between an Aβ_1-40_ fibril (hexamer or trimer) and a DPPC bilayer containing either no cholesterol, 30% cholesterol or 50 % cholesterol. Our simulation trajectories captured spontaneous binding of aqueous phase Aβ_1-40_ fibrils with the membranes and showed that 50% cholesterol leads to a significant increase in binding events. In this study, we observed that hydrophobic residues at the C-terminus (Ile31, Ile32, Met35, Val36, Val39, and Val40) and in the middle region (Leu17, Val18, Phe19, Phe20) play a key role in the binding of the Aβ_1-40_ fibrils with the membrane. This is in line with a previous study reporting that the above mentioned hydrophobic residues are most suitable for binding with lipid surfaces. Due to their hydrophobic nature, lipid surfaces accommodate the binding of such residues to reduce, to the extent possible, the exposure of their acyl chains to water [37]. Lys16, which is located next to the central hydrophobic cluster (residues 17-20) has also been previously reported to interact with the cell membrane [38]. Indeed, these data are in agreement with the previous CG study of Dias *et al*., which showed spontaneous binding of Aβ_1-42_ fibrils with membrane [21]. Our simulations reveal that both charged and hydrophobic residues participate in the binding with the membrane, in agreement also with previous studies reporting that both basic and hydrophobic amino acid residues play an important role in peripheral protein binding with the membranes [39, 40]. Furthermore, we noticed that C-terminal residues significantly participate in membrane binding, which is also in line with previous studies. For instance, Khemtemourian *et. al*. [41] showed that C-terminal hydrophobic residues of islet amyloid polypeptide (IAPP) can be inserted in the membrane, while Antonschmidt *et. al*. [42] showed that α-Synuclein fibrils contain a membrane binding domain at their C-terminal edge. We also observed a loss of water molecules and an increase in the number of lipid molecules in the vicinity of the Aβ_1-40_ fibrils upon membrane binding.

## 5. Conclusions

In summary, our simulations have shed light on the mechanism of Aβ1-40 fibrils interaction with the membrane. To the best of our knowledge, this is the first report where spontaneous binding of Aβ_1-40_ fibrils with a cholesterol-rich DPPC bilayer is shown. We identified key binding residues that stabilize the interaction of the Aβ1-40 fibrils with the membrane. These findings could guide the design of new drug candidates that could inhibit the association of Aβ1-40 fibrils with the membrane, thus opening new avenues in the design of inhibitors of the Aβ peptide-lipid membrane interaction.

## Acknowledgments

N.A. acknowledges the ERDF postdoc grant (No. 1.1.1.2/VIAA/4/20/757) for project funding. E.P. thanks the ERDF project BioDrug (No. 1.1.1.5/19/A/004) and the Latvian Council of Science (grant No. lzp-2020/2-0013) for financial support. We would like to thank Latvian Institute of Organic Synthesis for computational resources. N.A. would also like to thank the Centre for High Performance Computing (CHPC) in Cape Town (South Africa) for supercomputing resources.

## References

[1] C.M. Dobson, T.P. Knowles, M. Vendruscolo, The amyloid phenomenon and its significance in biology and medicine, Cold Spring Harbor perspectives in biology, 12 (2020) a033878.

[2] C.M. Dobson, The amyloid phenomenon and its links with human disease, Cold Spring Harbor perspectives in biology, 9 (2017) a023648.

[3] F. Chiti, C.M. Dobson, Protein misfolding, amyloid formation, and human disease: a summary of progress over the last decade, Annu. Rev. Biochem, 86 (2017) 27–68.

[4] N. Agrawal, E. Parisini, Early stages of misfolding of PAP248-286 at two different pH values: An insight from molecular dynamics simulations, Computational and Structural Biotechnology Journal, 20 (2022) 4892–4901.

[5] N. Agrawal, A.A. Skelton, Structure and function of Alzheimer’s amyloid βeta proteins from monomer to fibrils: a mini review, The protein journal, 38 (2019) 425–434.

[6] N. Agrawal, A.A. Skelton, 12-crown-4 ether disrupts the patient brain-derived amyloid-β-fibril trimer: Insight from all-atom molecular dynamics simulations, ACS chemical neuroscience, 7 (2016) 1433–1441.

[7] N. Agrawal, A.A. Skelton, Binding of 12-crown-4 with Alzheimer’s Aβ40 and Aβ42 monomers and its effect on their conformation: insight from molecular dynamics simulations, Molecular pharmaceutics, 15 (2017) 289–299.

[8] I.Y. Quiroga, A.E. Cruikshank, M. Bond, K. Reed, B.A. Evangelista, J.-H. Tseng, J.V. Ragusa, R.B. Meeker, H. Won, S. Cohen, Synthetic amyloid beta does not induce a robust transcriptional response in innate immune cell culture systems, Journal of neuroinflammation, 19 (2022) 1–12.

[9] A.-M. Chiorcea-Paquim, A.M. Oliveira-Brett, Amyloid beta peptides electrochemistry: A review, Current Opinion in Electrochemistry, 31 (2022) 100837.

[10] T. Wu, D. Lin, Y. Cheng, S. Jiang, M.W. Riaz, N. Fu, C. Mou, M. Ye, Y. Zheng, Amyloid Cascade Hypothesis for the Treatment of Alzheimer’s Disease: Progress and Challenges, AGING AND DISEASE, (2022).

[11] A. Mullard, Alzheimer amyloid hypothesis lives on, Nature Reviews Drug Discovery, 16 (2017) 3–5.

[12] H. Hampel, J. Hardy, K. Blennow, C. Chen, G. Perry, S.H. Kim, V.L. Villemagne, P. Aisen, M. Vendruscolo, T. Iwatsubo, The amyloid-β pathway in Alzheimer’s disease, Molecular Psychiatry, 26 (2021) 5481–5503.

[13] M.P. Murphy, H. LeVine iii, Alzheimer’s disease and the amyloid-β peptide, Journal of Alzheimer’s disease, 19 (2010) 311–323.

[14] S. Han, M. Kollmer, D. Markx, S. Claus, P. Walther, M. Fändrich, Amyloid plaque structure and cell surface interactions of β-amyloid fibrils revealed by electron tomography, Scientific Reports, 7 (2017) 43577.

[15] J. Kiskis, H. Fink, L. Nyberg, J. Thyr, J.-Y. Li, A. Enejder, Plaque-associated lipids in Alzheimer’s diseased brain tissue visualized by nonlinear microscopy, Scientific reports, 5 (2015) 13489.

[16] M.P. Burns, W.J. Noble, V. Olm, K. Gaynor, E. Casey, J. LaFrancois, L. Wang, K. Duff, Co-localization of cholesterol, apolipoprotein E and fibrillar Aβ in amyloid plaques, Molecular Brain Research, 110 (2003) 119–125.

[17] X. Yu, Q. Wang, Q. Pan, F. Zhou, J. Zheng, Molecular interactions of Alzheimer amyloid-β oligomers with neutral and negatively charged lipid bilayers, Physical Chemistry Chemical Physics, 15 (2013) 8878–8889.

[18] F. Tofoleanu, N.-V. Buchete, Molecular interactions of Alzheimer’s Aβ protofilaments with lipid membranes, Journal of molecular biology, 421 (2012) 572–586.

[19] F. Tofoleanu, B.R. Brooks, N.-V. Buchete, Modulation of Alzheimer’s Aβ protofilament-membrane interactions by lipid headgroups, ACS chemical neuroscience, 6 (2015) 446–455.

[20] X. Dong, Y. Sun, G. Wei, R. Nussinov, B. Ma, Binding of protofibrillar Aβ trimers to lipid bilayer surface enhances Aβ structural stability and causes membrane thinning, Physical Chemistry Chemical Physics, 19 (2017) 27556–27569.

[21] C.L. Dias, S. Jalali, Y. Yang, L. Cruz, Role of Cholesterol on Binding of Amyloid Fibrils to Lipid Bilayers, The Journal of Physical Chemistry B, 124 (2020) 3036–3042.

[22] M. Fändrich, S. Nyström, K.P.R. Nilsson, A. Böckmann, H. LeVine iii, P. Hammarström, Amyloid fibril polymorphism: a challenge for molecular imaging and therapy, Journal of Internal Medicine, 283 (2018) 218–237.

[23] J. Meinhardt, C. Sachse, P. Hortschansky, N. Grigorieff, M. Fändrich, Aβ (1-40) fibril polymorphism implies diverse interaction patterns in amyloid fibrils, Journal of molecular biology, 386 (2009) 869–877.

[24] A.K. Paravastu, R.D. Leapman, W.-M. Yau, R. Tycko, Molecular structural basis for polymorphism in Alzheimer’s β-amyloid fibrils, Proceedings of the National Academy of Sciences, 105 (2008) 18349–18354.

[25] U. Ghosh, K.R. Thurber, W.-M. Yau, R. Tycko, Molecular structure of a prevalent amyloid-β fibril polymorph from Alzheimer’s disease brain tissue, Proceedings of the National Academy of Sciences, 118 (2021) e2023089118.

[26] S. Jo, T. Kim, V.G. Iyer, W. Im, CHARMM-GUI: a web-based graphical user interface for CHARMM, Journal of computational chemistry, 29 (2008) 1859–1865.

[27] Y. Qi, H.I. Ingólfsson, X. Cheng, J. Lee, S.J. Marrink, W. Im, CHARMM-GUI martini maker for coarse-grained simulations with the martini force field, Journal of chemical theory and computation, 11 (2015) 4486–4494.

[28] D.H. de Jong, G. Singh, W.D. Bennett, C. Arnarez, T.A. Wassenaar, L.V. Schafer, X. Periole, D.P. Tieleman, S.J. Marrink, Improved parameters for the martini coarse-grained protein force field, Journal of chemical theory and computation, 9 (2013) 687–697.

[29] S.J. Marrink, H.J. Risselada, S. Yefimov, D.P. Tieleman, A.H. De Vries, The MARTINI force field: coarse grained model for biomolecular simulations, The journal of physical chemistry B, 111 (2007) 7812–7824.

[30] M. Bixon, S. Lifson, Potential functions and conformations in cycloalkanes, Tetrahedron, 23 (1967) 769–784.

[31] M. Parrinello, A. Rahman, Polymorphic transitions in single crystals: A new molecular dynamics method, Journal of Applied physics, 52 (1981) 7182–7190.

[32] G. Bussi, D. Donadio, M. Parrinello, Canonical sampling through velocity rescaling, The Journal of chemical physics, 126 (2007) 014101.

[33] M.J. Abraham, T. Murtola, R. Schulz, S. Páll, J.C. Smith, B. Hess, E. Lindahl, GROMACS: High performance molecular simulations through multi-level parallelism from laptops to supercomputers, SoftwareX, 1 (2015) 19–25.

[34] S. Buchoux, FATSLiM: a fast and robust software to analyze MD simulations of membranes, Bioinformatics, 33 (2017) 133–134.

[35] P. Bharadwaj, T. Solomon, C.J. Malajczuk, R.L. Mancera, M. Howard, D.W. Arrigan, P. Newsholme, R.N. Martins, Role of the cell membrane interface in modulating production and uptake of Alzheimer’s beta amyloid protein, Biochimica et Biophysica Acta (BBA)-Biomembranes, 1860 (2018) 1639–1651.

[36] S.M. Butterfield, H.A. Lashuel, Amyloidogenic protein–membrane interactions: mechanistic insight from model systems, Angewandte Chemie International Edition, 49 (2010) 5628–5654.

[37] A.K. Srivastava, J.M. Pittman, J. Zerweck, B.S. Venkata, P.C. Moore, J.R. Sachleben, S.C. Meredith, β-Amyloid aggregation and heterogeneous nucleation, Protein Science, 28 (2019) 1567–1581.

[38] S. Sinha, D.H. Lopes, G. Bitan, A key role for lysine residues in amyloid β-protein folding, assembly, and toxicity, ACS Chemical Neuroscience, 3 (2012) 473–481.

[39] E. Fuglebakk, N. Reuter, A model for hydrophobic protrusions on peripheral membrane proteins, PLoS computational biology, 14 (2018) e1006325.

[40] A.H. Larsen, L.H. John, M.S. Sansom, R.A. Corey, Specific interactions of peripheral membrane proteins with lipids: what can molecular simulations show us?, Bioscience Reports, 42 (2022) BSR20211406.

[41] L. Khemtemourian, H. Fatafta, B. Davion, S. Lecomte, S. Castano, B. Strodel, Structural dissection of the first events following membrane binding of the islet amyloid polypeptide, Frontiers in molecular biosciences, 9 (2022) 849979.

[42] L. Antonschmidt, R. Dervişoğlu, V. Sant, K. Tekwani Movellan, I. Mey, D. Riedel, C. Steinem, S. Becker, L.B. Andreas, C. Griesinger, Insights into the molecular mechanism of amyloid filament formation: Segmental folding of α-synuclein on lipid membranes, Science Advances, 7 (2021) eabg2174.

